# Simultaneous expression of UV and violet SWS1 opsins expands the visual palette in a group of freshwater snakes

**DOI:** 10.1101/2021.09.07.459283

**Authors:** Einat Hauzman, Michele E.R. Pierotti, Nihar Bhattacharyya, Juliana H. Tashiro, Carola A.M. Yovanovich, Pollyanna F. Campos, Dora F. Ventura, Belinda S.W. Chang

## Abstract

Snakes are known to express a rod visual opsin and two cone opsins, only (SWS1, LWS), a reduced palette resulting from their supposedly fossorial origins. Dipsadid snakes in the genus *Helicops* are highly visual predators that successfully invaded freshwater habitats from ancestral terrestrial-only habitats. Here we report the first case of multiple SWS1 visual pigments in a vertebrate, simultaneously expressed in different photoreceptors and conferring both UV and violet sensitivity to *Helicops* snakes. Molecular analysis and *in vitro* expression confirmed the presence of two functional SWS1 opsins, likely the result of recent gene duplication. Evolutionary analyses indicate that each *sws1* variant has undergone different evolutionary paths, with strong purifying selection acting on the UV-sensitive copy and d_N_/d_S_ ∼1 on the violet-sensitive copy. Site-directed mutagenesis points to the functional role of a single amino acid substitution, Phe86Val, in the large spectral shift between UV and violet opsins. In addition, higher densities of photoreceptors and SWS1 cones in the ventral retina suggest improved acuity in the upper visual field possibly correlated with visually-guided behaviors. The expanded visual opsin repertoire and specialized retinal architecture are likely to improve photon uptake in underwater and terrestrial environments, and provide the neural substrate for a gain in chromatic discrimination, potentially conferring unique color vision in the UV-violet range. Our findings highlight the innovative solutions undertaken by a highly specialized lineage to tackle the challenges imposed by the invasion of novel photic environments and the extraordinary diversity of evolutionary trajectories taken by visual opsin-based perception in vertebrates.

## Introduction

The visual system of extant snakes is thought to be, at least in part, the result of an ancestral fossorial and nocturnal habitat (Walls 1942; Hsiang et al. 2015), resulting in the loss of many visual structures, including ocular ciliary muscles and scleral ossicles (Walls 1942; Ott 2006), photoreceptor types and organelles (e.g. double cones, oil droplets), and genes involved in visual processing, including two of the four vertebrate cone opsin classes (SWS2 and RH2), otherwise present in the lizards (Walls 1942; Underwood 1967; Davies et al. 2009; Simões et al. 2015; Emerling 2017). While the nature of the ancestral snake habitat is still debated (e.g. Hsiang et al. 2015; Caprette et al. 2004; Lee et al. 2016), the unique origin of novel features in response to diurnality, such as an exclusive type of double cone (Walls 1942), cone microdroplets (Wong 1989) and transmuted photoreceptors (Walls, 1942), is consistent with such an evolutionary transition. In particular, the unique phenomenon of transmutation whereby the typical morphological and functional distinction between vertebrate cones and rods leaves way to transitional states and physiological interconversion between cones and rods (cone-like rods and rod-like cones) (Walls 1942; Simões et al. 2015; Schott et al. 2016; Bhattacharyya et al. 2017; Hauzman et al. 2017) suggests the evolution of novel solutions in response to constraints (gene loss) brought about by past evolutionary history.

The visual processing begins with the absorption of photons by a light-sensitive derivative of vitamin A, the chromophore, covalently bound to a G protein-coupled receptor, the opsin. Photoisomerization of the chromophore induces a conformational change in the opsin initiating signal transduction in the cone and rod photoreceptor cells (Burns and Lamb 2004). The type of chromophore used (either vitamin A1- or vitamin A2-based) and the amino acid sequence of the opsin molecule determine its spectral absorption peak (*λ*_max_), with substitutions at key amino acid residues shifting the *λ*_max_ to shorter or longer wavelengths (Yokoyama 2002). Snakes are thought to use only A1-derivative retinal, based on recorded *λ*_max_ from microspectrophotometry (MSP) (e.g. Sillman et al. 1997; Davies et al. 2009; Hart et al. 2012) or HPLC analysis (Seiko et al. 2020), and exhibit a high number of evolutionary changes in their (reduced) opsin gene set, including substitutions at functional sites unique to this clade of reptiles (Simões et al. 2016).

The henophidian snakes, a non-monophyletic group of largely nocturnal species, are thought to exhibit a retina with both cones and rods photoreceptors (duplex), possibly reflecting the ancestral snake condition, with a high density of rods containing the rod-typical dim light photopigment (rhodopsin, RH1), and two types of cones with photopigments sensitive to short (SWS1) and long (LWS) wavelengths (Davies et al. 2009). In the derived monophyletic clade of caenophidian (“advanced”) snakes, nocturnal species possess duplex retinas (Walls 1942; Hauzman et al. 2017). Diurnal species, however, exhibit “all-cone” retinas in which transmuted cone-like rods exhibit a cone-like gross morphology, with shorter outer segments, but with a rod ultrastructure and photopigment (RH1) (Schott et al. 2016; Bhattacharyya et al. 2017), and lower overall density of photoreceptors (Hauzman et al. 2017). Frequently, in these retinas, the rhodopsin has a considerable blue shift in *λ*_max_ from the typical ∼495-500nm of most vertebrates (Yokoyama 2000), to ∼484nm (Sillman et al. 1997; Schott et al. 2016; Simões et al. 2016; Bhattacharyya et al. 2017; but see Hart et al. 2012; Simões et al. 2020; Seiko et al. 2020). These drastic changes in retinal morphology and rhodopsin function seem to be associated with a gain in visual acuity and a possible increase in chromatic discrimination (Schott et al. 2016).

Adaptation to different light environments brings about habitat-specific demands on the species’ visual system. In particular, transitions to aquatic environments impose considerable evolutionary pressures on the opsin genes, due to the great photic variability, reduction of available light, and filtering of certain wavelengths through the water column (Loew and Lythgoe 1985; Lythgoe and Partridge 1989; Bowmaker and Hunt 2006). In marine elapid snakes, for instance, blue-shifts of the long-wavelength opsins in deep-diving species might represent adaptations to the available light field in deeper waters (Seiko et al. 2020), while spectral shifts of the SWS1 opsins from the ancestral UV toward longer wavelengths might improve photon capture in marine waters (Hart et al. 2012). In the marine genus *Hydrophis* (Elapidae), a recent study described the presence of allelic polymorphisms, across multiple species, at residue 86 (Simões et al. 2020), known to cause major shifts from UV to violet sensitivity in non-bird vertebrates (Hunt et al. 2007). Since genomic data in at least two species showed the presence of individuals polymorphic for SWS1 alleles, the authors speculated that both alleles might be simultaneously expressed, either in distinct visual cells or coexpressed in the same photoreceptor, potentially expanding the breadth of sensitivity in the UV-violet range. Similarly, in a genus of freshwater dipsadid snakes, *Helicops*, two studies found polymorphisms at the same SWS1 residue (Phe/Val86) (Simões et al. 2016; Hauzman et al. 2017) and suggested a possible functional advantage associated with improved sensitivity in aquatic environments. Nevertheless, in both these independent lineages of aquatic snakes, it is, at present, not known whether both alleles are expressed in the same individual. In addition, if such condition were to be demonstrated, its functional importance would depend on their site of expression within the outer retina: if both opsins were simultaneously expressed in different photoreceptors, they would give rise to distinct sensitivity curves in the UV-violet range with the potential for expanded discrimination in the UV-violet range, but if they were coexpressed within the same cone cell, they would simply give rise to a single photoreceptor class intermediate in wavelength sensitivity between the Phe86 and the Val86 SWS1 opsin. Finally, we lack a functional characterization of the Phe/Val86 substitution and its effects on SWS1 wavelength sensitivity in any snake species.

The aquatic *Helicops* snakes occupy a variety of lotic and lentic freshwater environments from very clear to murky waters (Lema et al. 1983; Martins and Oliveira 1998; De Aguiar and Di-Bernardo 2004). Freshwater habitats have the most variable and complex underwater light fields, where dissolved organic and inorganic matter affect scatter and the selective filtering of wavelengths and where depth and the nature of the substrate (e.g. sand, clay, rocks, vegetation) can lead to rapid changes in the characteristics of the photic environment at both temporal and spatial scale (Loew and Lythgoe 1985; Lythgoe and Partridge 1989), imposing great challenges to the animals’ visual system. *Helicops* species perform most of their activities and spend most of their time in the water, including actively hunting (Martins and Oliveira 1998; De Aguiar and Di-Bernardo 2004; Ávila et al. 2006), resting and reproduction (Martins and Oliveira 1998; Ávila et al. 2006). Predominantly nocturnal, they use the visual sense to hunt fish at the surface, in the water column, or at the bottom, and even venturing on land to prey on anurans near water banks (Martins and Oliveira 1998; De Aguiar and Di-Bernardo 2004). Compared to terrestrial relatives, *Helicops* snakes have eyes and nostrils more dorsally positioned in the head, a morphological adaptation to an aquatic lifestyle (Scartozzoni 2005), allowing breathing and viewing above the water surface while the body remains submerged. These behaviors seem to underscore an exceptional versatility of *Helicops*’ visual system.

In this study, we applied a range of experimental and computational approaches to investigate how the invasion of *Helicops* snakes into diverse, complex freshwater habitats shaped the evolution of their visual system in terms of retinal structure, spectral sensitivity, and opsin gene complement. We find that *Helicops* snakes have up to four visual pigments simultaneously expressed in their retinas, with two distinct SWS1 opsins conferring sensitivity in the UV and the violet spectrum, the first documented case of UV + violet sensitivity based on two SWS1s in a vertebrate species. Phylogenetic reconstruction, sequence divergence, and selection analysis support a recent gene duplication, with evidence for positive selection at the ancestral branch leading to *Helicops*, and distinct selection trajectories for the two *sws1* paralogs. Using site-directed mutagenesis, we demonstrate that a single amino acid substitution is responsible for the amplitude of spectral shift between the UV and violet opsin. Morphological analyses reveal a specialized retinal architecture (e.g. a ventral *area centralis*) associated with specific visually-guided behaviors, such as diving to escape from aerial predators. Our results indicate that these aquatic hunter snakes that operate at the air/water interface have evolved a unique solution among vertebrates to improve photon capture over a broad range of wavelengths at the short end of the visual spectrum, with the simultaneous expression of two functionally distinct SWS1 opsins, a configuration that might even afford color vision in the UV-violet, with exciting implications for sensory perception in this group and, in general, for the evolution of color vision in vertebrates.

## Results

### Four visual opsins are simultaneously expressed in the retinas of *Helicops* snakes

We sequenced PCR products of almost the entire coding region (∼1,000 bp) of the visual opsin genes, *sws1*, *lws*, and *rh1*, expressed in the retinas of single specimens from three *Helicops* species, *H. infrataeniatus*, *H. carinicaudus*, and *H. leopardinus*, and a closely related aquatic species *Hydrops caesurus*. Identities of the opsins genes were confirmed using BLAST searches and phylogenetic analysis (supplementary figs. S1 and S2). For the *sws1* gene, we sequenced multiple individually selected (white/blue screened) positive clones (∼30) of the four species mentioned above and, additionally, of *Helicops modestus*. Analysis of the sequences revealed the presence of two distinct *sws1* variants expressed in retinas of *H. infrataeniatus*, *H. carinicaudus*, and *H. modestus*, hereafter indicated as *sws1a* and *sws1b* (fig. 1a; supplementary fig. S2). A maximum likelihood phylogenetic reconstruction grouped the *Helicops sws1b* sequences in a monophyletic clade with high bootstrap support and nested within a paraphyletic *sws1a* branch (fig. 1a; supplementary fig. S2). The *sws1a* and *sws1b* sequences of each species had ∼98% of similarity, a level comparable to that of the paralogous *lws* and *mws* opsin genes of Old World monkeys (Nathans et al. 1986). The *sws1* variants had a considerable amount of non-synonymous substitutions (up to 5% of the residues), including two important spectral tuning sites, 86 and 93 (supplementary fig. S3). In the three *Helicops* species, *H. infrataeniatus*, *H. carinicaudus*, and *H. modestus*, the SWS1A opsin has Phe86 which indicates sensitivity in the UV range (Cowing et al. 2002; Fasick et al. 2002; Parry et al. 2004), while the SWS1B opsin has a Phe86Val substitution, and thus, a sensitivity predicted in the violet range (Parry et al. 2004) (supplementary fig. S3 and table S1). In one species, *H. leopardinus*, the amplified cDNA sequences obtained from a single retina of one specimen revealed only one *sws1* variant, identified as *sws1a* based on phylogenetic reconstruction (fig. 1a; supplementary fig. S2), and with residue Phe86 (supplementary fig. S3 and table S1), conferring UV sensitivity. The counter-eye of the same *H. leopardinus* individual was analyzed with microspectrophotometry revealing the presence of a violet pigment (see results in the following section), and indicating that both *sws1* variants are expressed in the same individual.

**Figure 1.**
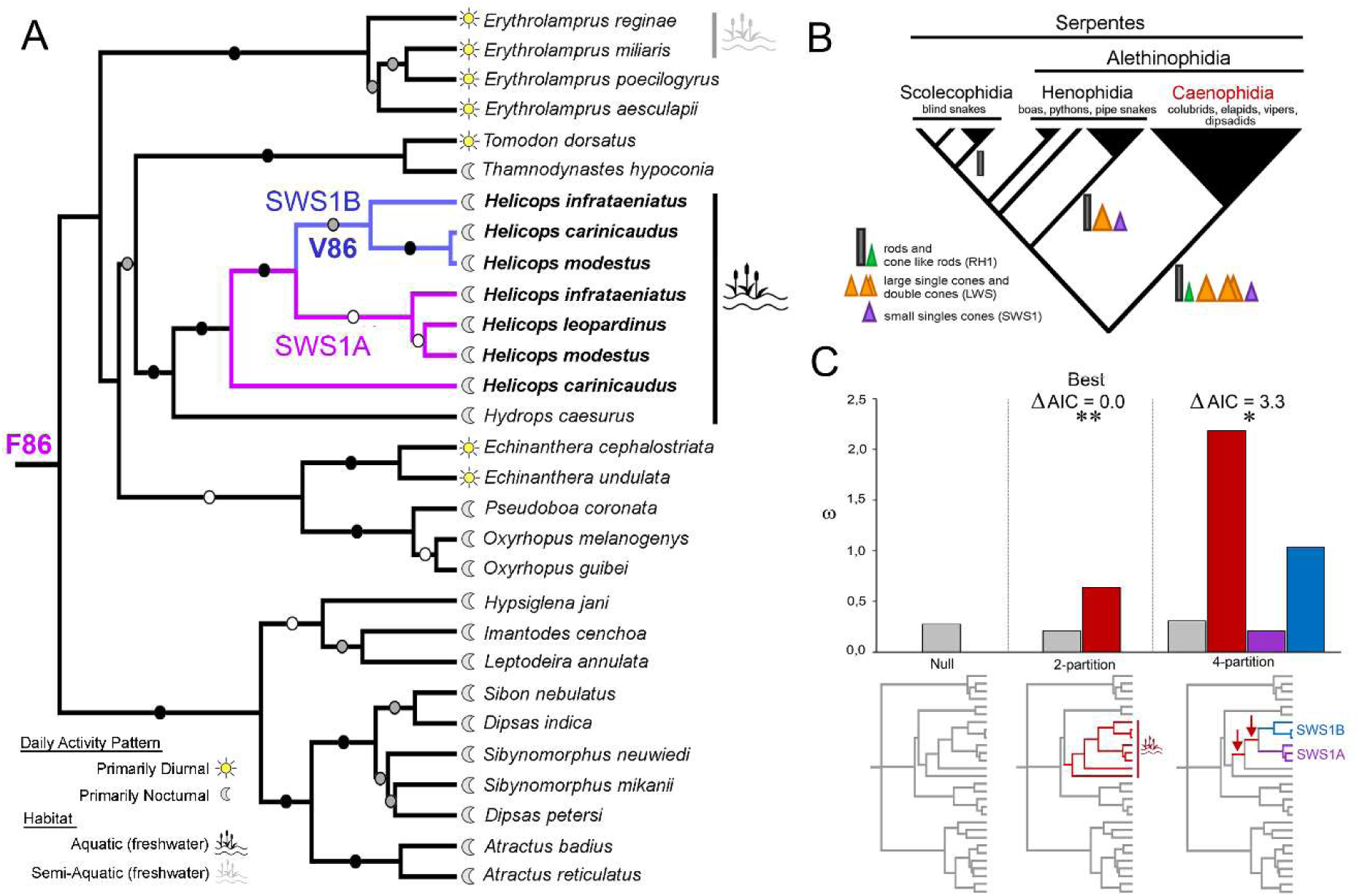
Dipsadidae sws1 gene tree and patterns of molecular evolution. (A) Dipsadidae *sws1* gene tree. Aquatic and semi-aquatic lineages are indicated, and diel activity patterns are depicted based on the literature (Martins and Oliveira 1998; Torello-Viera and Marques 2017). *Helicops* species are highlighted in bold, and *sws1a* and *sws1b* branches are differentiated by purple and blue lines, respectively. Maximum Likelihood (ML) bootstrap supports are represented for each resolved node by black (86-100%), gray (71-85%), and white (56–70%) circles. (B) Schematic cladogram showing the relationships between the major groups of snakes and photoreceptors. (C) Comparison of divergent omega classes (ω) among partitions obtained in CmC analysis, and the respective ω2 from the null model. Topologies below x-axis represent the partitions used in CmC. Lineages highlighted on each tree were included as foreground clade, and are differentiated by the colors in the tree; lineages in grey represent the background clade. The following partition models are shown: a two-partition isolating aquatic snakes as foreground, and a four-partition isolating the *Helicops* ancestral branches (arrows), the *sws1a* and the *sws1b*. LRTs with Χ^2^ distribution were performed to compare CmC with the null model. Statistical significance is indicated by *p < 0.02 and **p < 0.002. The likelihood of each partition model was compared using differences in AIC. The partitions that best fit the data is indicated.

Additionally, we searched for the presence of more than one *sws1* variant by sequencing the exon 1 from genomic DNA of single individuals of the *Helicops* species: *H. angulatus*, *H. gomesi*, *H. hagmanni*, *H. leopardinus*, and *H. polylepis*, and other aquatic dipsadids, *Hydrops triangularis*, *Hydrops martii*, *Pseudoeryx plicatilis*, *Sordellina punctata*, and *Hydrodynastes gigas*. In all *Helicops* species, we consistently found both residues Phe86 and Val86 (supplementary fig. S4, Supplementary Material online), indicating that the presence of two short-wavelength sensitive opsins, a UV and a violet, within individuals, is widespread in the genus. In all other species (non-*Helicops*) investigated, we found the presence of residue Phe86, only, indicating a single UV-sensitive SWS1 opsin.

Furthermore, we analyzed intronic sequences of single individuals from six *Helicops* species. We amplified by PCR and sequenced the intron 1 of *H. angulatus*, *H. gomesi, H. hagmanni*, *H. leopardinus*, *H. modestus*, and *H. polylepis*. The amplified fragments were isolated from agarose gel (supplementary fig. S5) and sequenced in both directions. All species had an intron 1 with approximately 950 bp, except *H. gomesi*, with ∼1555 bp (Supplementary Material online). In individuals of two species, *H. modestus* and *H. leopardinus*, we amplified and sequenced a second and highly length-divergent intron 1, with 468 bp (supplementary fig. S5, Supplementary Material online). This finding strongly indicates that the two *sws1* variants represent distinct paralogous genes. Moreover, based on the electrophoresis images (supplementary fig. S5) and preliminary sequencing analyses (data not shown), we identified signals of the presence of three distinct intron 1s with different lengths in *H. angulatus* and *H. modestus*, which suggests the possibility of multiple gene copies and deserves further molecular analyses in future studies.

### UV and violet opsins of *Helicops* evolved under different patterns of selection

We searched for distinct signatures of selection acting on the *sws1* opsin genes of *Helicops* snakes (fig. 1c) by estimating the ratio of nonsynonymous to synonymous substitutions (d_N_/d_S_ or ω) in a Dipsadidae alignment, using random-site, branch-site, and clade models (CmC) with PAML. The analysis based on CmC provided evidence for positive selection at the ancestral branch leading to *Helicops* (ω = 2.2) (fig. 1c), consistent with accelerated rates of evolution after a *sws1* gene duplication in the *Helicops* lineage. In a four-partition CmC model, we found signals of strong purifying selection on the *sws1a* clade (ω = 0.2), but higher rates of d_N_/d_S_ on the *sws1b* clade, with ω = 1.0. Using CmC models, we also tested partitions isolating as foregrounds, respectively: 1) the aquatic lineage (Hydropsini snakes represented here by *Helicops* species and *Hydrops caesurus*), and 2) the *Helicops* clade. Both foreground clades had higher rates of d_N_/d_S_ compared to the background (fig. 1c; supplementary fig. S6 and table S2). In random-site models, the average ω value estimated under the null model (M0) was 0.11, and significant rates of substitution were variable across sites (M3 vs. M0), as expected for a protein-coding gene under strong purifying selection. However, no evidence of positive selection (ω > 1) on the *sws1* was detected along the Dipsadidae alignment (M2a vs. M1a, M8 vs. M7; supplementary table S3). We did not find evidence for positive selected sites in specific clades of interest (aquatic snakes, *Helicops, sws1a*, and *sws1b*), using branch-site models (supplementary table S4).

### Functional assessment of the visual opsins of *Helicops* snakes

We used micro-spectrophotometry (MSP) of retinas on 2 wild-caught *H. leopardinus* and 1 wild-caught *H. angulatus* for an *in-situ* assessment of the visual pigments in these snakes. Given the limited number of records, we did not attempt accurate characterizations of peak absorbance values, however, these preliminary data were useful to provide a first qualitative evaluation of the theoretical *λ*_max_ predictions and confirm the presence of distinct UV/violet photoreceptor classes within a single retina. In the *H. angulatus* individual, we recorded a UV-sensitive SWS1 single cone, with *λ*_max_ at 395nm (n=1) and a violet-sensitive single cone with *λ*_max_ at 420.5±3.5nm (n=2) (fig. 2a; supplementary fig. S7 and table S5). In *H. leopardinus*, we recorded a UV-sensitive SWS1 single cone, with *λ*_max_ at 363nm (n=1) in one individual and a violet-sensitive single cone with *λ*_max_ at 411nm (n=1) in the second individual tested (fig. 2b; supplementary fig. S7 and table S5). The contralateral eye of this second individual was used for the genetic analysis revealing the expression of a UV (Phe86) *sws1a* opsin gene. The combined MSP and genetic results for this individual of *H. leopardinus* and the MSP results from the *H. angulatus* individual are an additional confirmation that UV and violet pigments can be both expressed simultaneously in the same individual, and in distinct photoreceptors.

**Figure 2.**
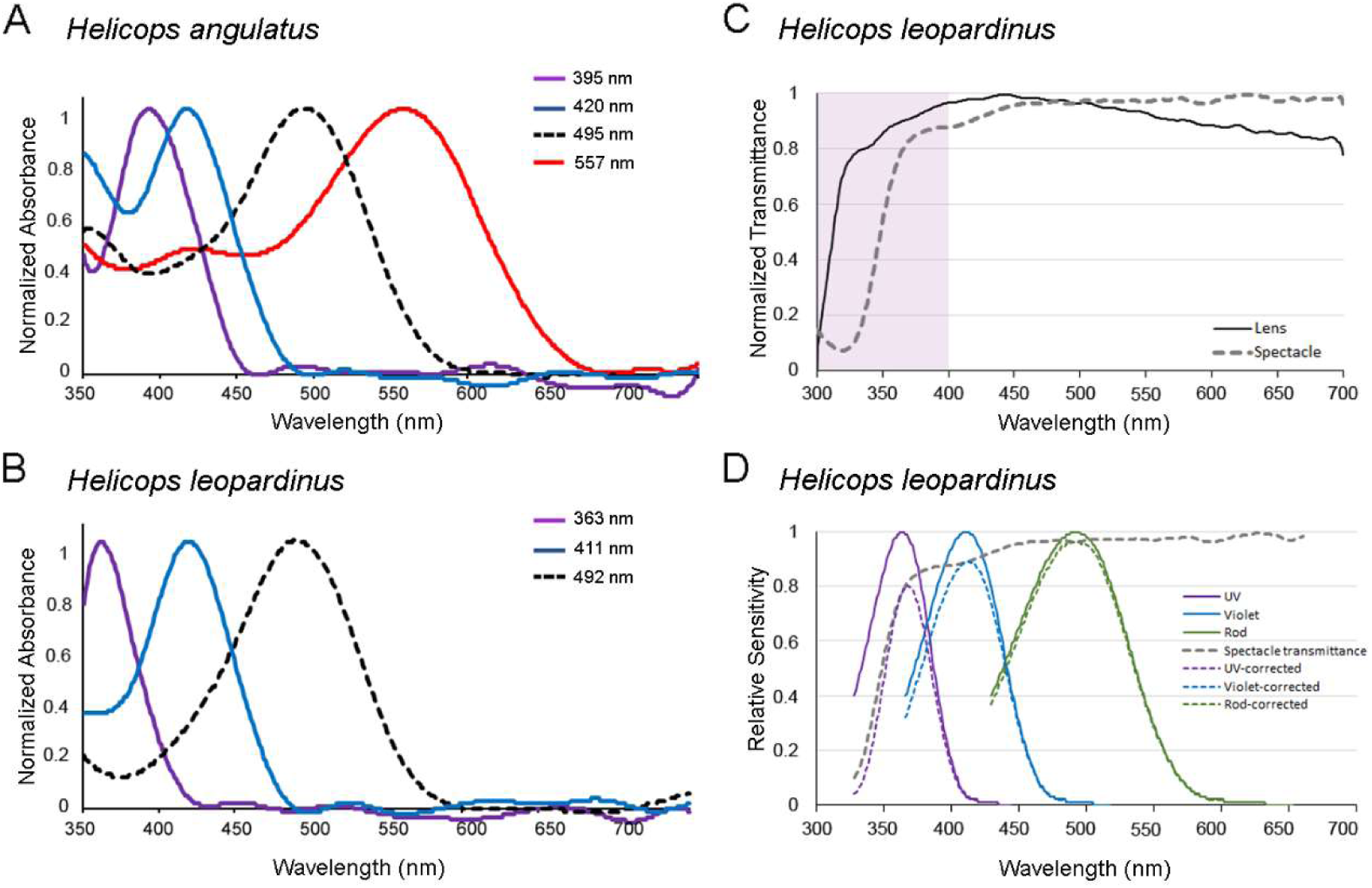
Four visual opsins are simultaneously expressed in retinas of *Helicops* snakes. (A, B) MSP measurements of *Helicops angulatus* and *Helicops leopardinus*: interpolating Gaussian fit to the raw MSP data (see also supplementay fig. S6 for representative MSP records). (C) Lens and spectacle normalized transmittance of *H. leopardinus*. (D) Effect of spectacle transmittance on spectral sensitivity in *H. leopardinus*: continuous lines represent photoreceptor spectral sensitivity templates as in (A), and dashed lines represent the same spectra corrected by the effect of the spectacle.

In *H. angulatu*s, we found rods with maximum absorbance at 495±2nm (n=11) (fig. 2a; supplementary fig. S7 and table S5) in full agreement with theoretical predictions based on the amino acids found in spectral tuning sites (Simões et al. 2016) (supplementary table S6), and an LWS visual pigment with *λ*_max_ at 557nm (n=1) (fig. 2a; supplementary fig. S7 and table S5), a value close to the predicted *λ*_max_ of ∼555nm determined based on the five sites rule (Yokoyama and Radlwimmer 1998), and the substitution Ser164Ala (Simões et al. 2016) (supplementary table S7). In *H. leopardinus,* however, rods had maximum absorbance at 493±2nm (n=8) (fig. 2b; supplementary fig. S7 and table S5), a value that is substantially different from our predictions based on molecular analysis (supplementary table S7), which revealed the amino acids Asn83, Ser292, and Ala299, known to cause a ∼16 nm blue-shift and generate a *λ*_max_ at ∼484nm. This amino acid combination and spectral peak were described in diurnal colubroids (Schott et al. 2016; Simões et al. 2016; Bhattacharyya et al. 2017; Hauzman et al. 2017), implying that in *H. leopardinus*, other residues might be involved in the rhodopsin spectral tuning.

Analysis of ocular media transmittance of two individuals of *H. leopardinus* showed a highly UV-A transmissive lens, with a 50% cut-off transmission (*λ*_T50_) at 312nm, and spectacle with *λ*_T50_ at 352nm (fig. 2c). A spectacle more UV-absorptive than the lens has been reported for other snake species (Simões et al. 2016). We calculated the impact of the ocular media transmittance in visual sensitivity and the amount of incoming light available to the visual system by quantifying the effect of the spectacle transmittance on photoreceptors spectral sensitivity curves of *H. leopardinus* (fig. 2d). The spectral absorption peak of the UV opsin was only marginally affected by the spectacle (+3nm) (fig. 2d).

### Molecular mechanisms underlying the spectral tuning of UV and violet opsins of *Helicops*

We investigated the spectral phenotype of *H. modestus* SWS1 opsins and the molecular mechanisms underlying their functional divergence using spectroscopy assays of wildtype and mutant SWS1 expressed and purified *in vitro*. *Sws1a* and *sws1b* of *H. modestus* were each individually ligated into the p1D4-hGFP II expression vector, which was then used for heterologous expression in HEK293T cells. The expressed proteins were purified by immunoaffinity and reconstituted with 11-*cis*-retinal chromophore in the dark. Spectroscopy of the purified proteins revealed that both visual pigments are functional and have distinct absorption peaks. The SWS1A, with the amino acids Phe86/Met93, produced a dark absorbance spectrum in the UV range, with *λ*_max_ at 363.5±2.4nm (n=12), and the SWS1B, with Val86/Val93, generated a dark absorbance spectrum in the violet range, with *λ*_max_ at 416.8±0.4nm (n=6) (fig. 3).

**Figure 3.**
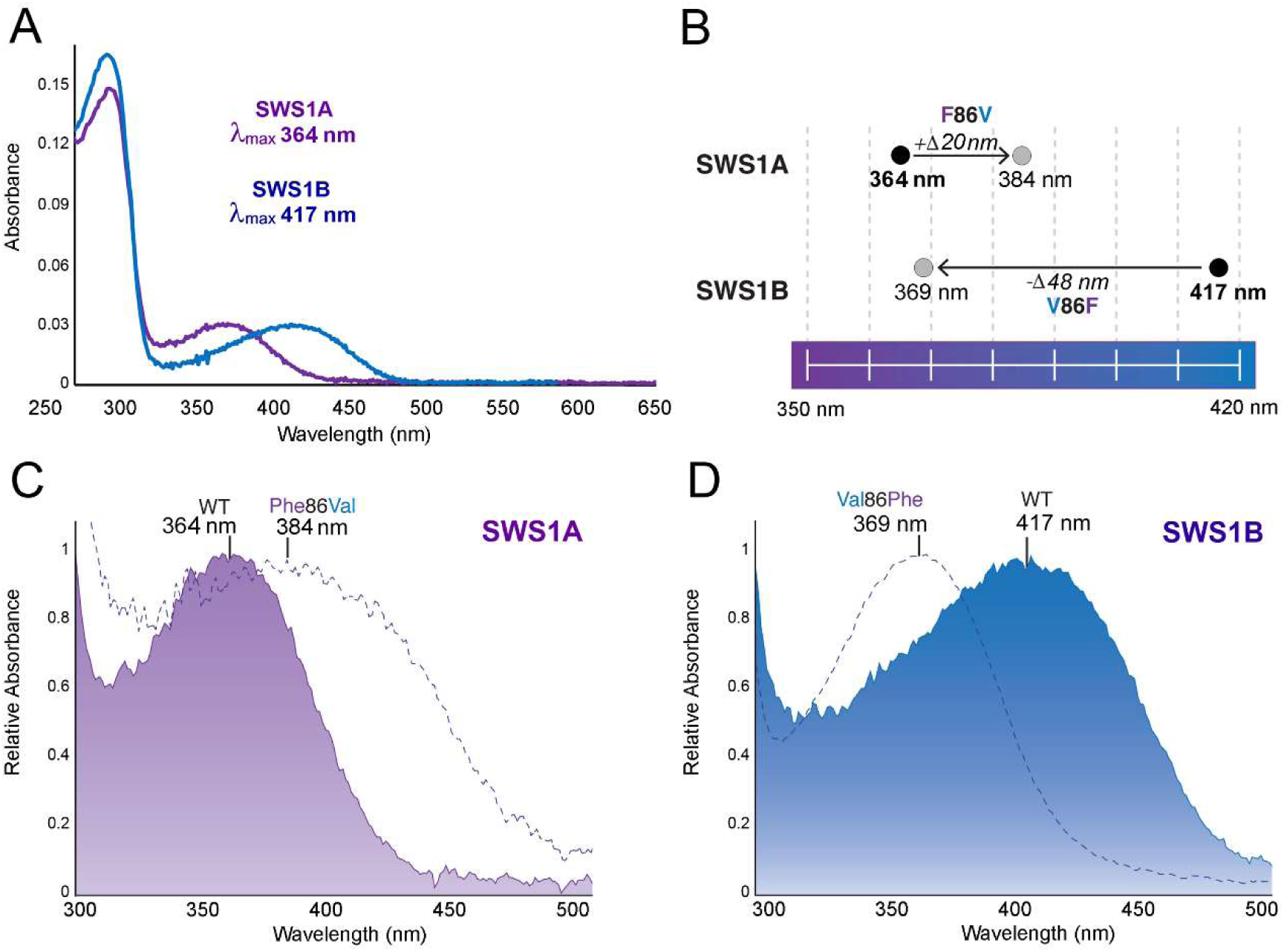
Molecular mechanisms underlying the spectral tuning of UV and violet opsins of *Helicops*: UV-visible dark absorption spectra of *H. modestus* SWS1 opsins reconstituted *in vitro*. (A) Dark spectra for wild SWS1A and SWS1B opsins. (B) Spectral differences between wild types and SWS1A and SWS1B mutants. (C) Spectral difference between wild SWS1A and the mutant Phe86Val. (D) Spectral difference between wild SWS1B and the mutant Val86Phe. λmax estimations are shown for each visual pigment. WT, wildtype.

We explored the functional role of the single amino acid substitution Phe86Val using site-directed mutagenesis and *in vitro* expression of mutant opsins. The substitution Phe86Val in an SWS1A background caused a ∼20nm shift towards the violet, with a spectral absorption peak at 384.1±0.6nm (n=4). Conversely, the substitution Val86Phe in an SWS1B background caused a ∼48nm shift, and generated a UV visual pigment, with *λ*_max_ at 369.0±0.1nm (n=3), only 4nm different from the wildtype SWS1A opsin (fig. 3). These results show experimentally, for the first time in snakes, that residue 86 is responsible for a major shift between UV and violet opsins, while other residues might have minor additional effects on spectral tuning. The substitution Val86Phe in a violet opsin background generated a complete shift toward the UV-band, indicating that Phe86 might cause the loss of electrostatic protonation of the chromophore Schiff base (Cowing et al. 2002; Parry et al. 2004). However, the spectral peak of the mutated UV opsin, Phe86Val, did not cause the complete shift toward the violet band, indicating that other residues present in the violet opsin (SWS1B) might be necessary for fully stabilizing the protonation of the Schiff base (Fahmy and Sakmar 1993), resulting in the difference in the degree of spectral shift in the forward and backward mutations.

### The retinal morphology of *Helicops* snakes has a pattern similar to diurnal colubroids

We investigated the photoreceptor types, densities, and distribution in retinas of *Helicops* snakes, using immunohistochemistry. *Helicops* snakes are predominantly nocturnal (Martins and Oliveira 1998; De Aguiar and Di-Bernardo 2004), yet retinal sections of *H. modestus* and *H. carinicaudus* showed a structure more typical of diurnal colubroids, with a single layer of photoreceptors *nuclei* in a thin outer nuclear layer, with about 10 µm thickness, indicating low photoreceptor density (fig. 4). Additionally, the absence of typical rods with long and slender outer segment is characteristic of primarily diurnal species (Walls 1942; Underwood 1967; Wong 1989; Sillman et al. 1997; Hauzman et al. 2014; Schott et al. 2016; Hauzman et al. 2017). Nocturnal colubroids, on the other hand, have duplex retinas, with a high amount of typical rods and a thick outer nuclear layer (Hauzman et al. 2017). Using different combinations of specific anti-opsin antibodies (double-labeling), previously used in snake retinas (Schott et al. 2016; Bhattacharyya et al. 2017; Hauzman et al. 2017; Bittencourt et al. 2019), we identified four photoreceptor types. The retinas are dominated by the LWS cone class with two subpopulations, large single cones and double cones (fig. 4). Small single SWS1 cones represent a small population of photoreceptors, and a fourth group of photoreceptors contain the rhodopsin (RH1) photopigment and were classified as cone-like rods (fig. 4). These had outer segments of comparable length of that observed for LWS and SWS1 cones, but with less bulbous inner segments, as previously described in the “all-cone” retinas of diurnal colubroids, in which the rhodopsin photopigment is expressed in a group of transmuted cone-like rods (Hauzman et al. 2014; Schott et al. 2016; Bhattacharyya et al. 2017; Hauzman et al. 2017).

**Figure 4.**
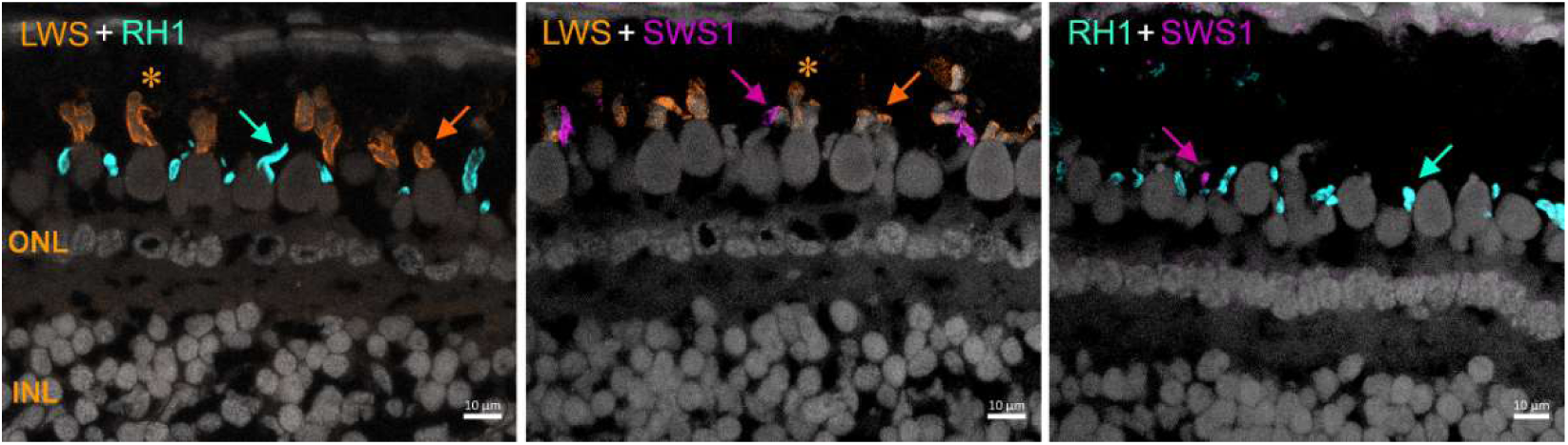
The retinal morphology of *Helicops*. Confocal images of retinal sections of *H. modestus* double-labeled with anti-opsins antibodies (arrows): LWS cones highlighted in orange, cone-like rods (RH1) in turquoise, and SWS1 cones in magenta. The asterisks (*) indicate double cones (LWS), with a large principal member and slender accessory member. No opsin colocalization was observed in any cone. Cell *nuclei* in the outer nuclear layer (ONL) and inner nuclear layer (INL) were stained by DAPI and are differentiated in gray.

We analyzed the total density and distribution of the photoreceptors and of the SWS1 cones in three whole-mounted retinas of *H. modestus* using a stereological approach (supplementary table S8). The total photoreceptor population estimated was 85,795 *±* 33,181 cells, with a mean density of 9,585 *±* 1,729 cells/mm_2_ (supplementary table S9), a low mean density value, similar to that described for diurnal dipsadids (Hauzman et al. 2014). The estimated population of SWS1 cones was 6,374 *±* 605 cells, with a mean density of 751 *±* 199 cells/mm_2_, and it accounted for 8% *±* 2.3 of the photoreceptor population (supplementary table S9). The topographic maps showed higher densities of total photoreceptor and of the SWS1 cones in the ventral retina, with anisotropic *area centralis* (fig. 5). This type of specialization is likely to provide higher visual acuity in the upper visual field and suggest a higher absorption of short-wavelength photons from above.

**Figure 5.**
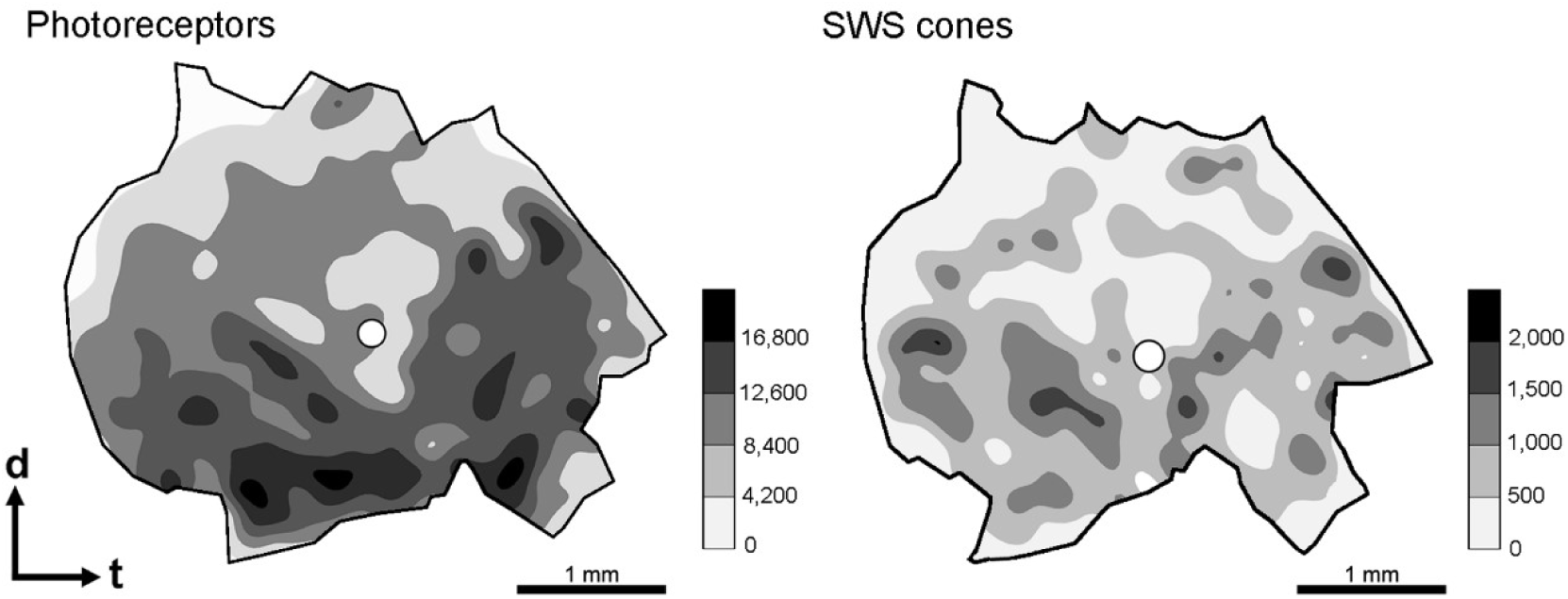
The retinal topographic maps of *Helicops modestus*. Total photoreceptors and SWS1 cones, with anisotropic *area centralis* in the ventral region. Gray bars represent the number of cells/mm^2^. The optic nerve head is depicted as a white circle. d, dorsal; t, temporal.

## Discussion

In this study we showed that two distinct SWS1 opsins with UV- or violet sensitivity can be expressed simultaneously in the retina of the aquatic dipsadid snake *Helicops* and that a single amino acid substitution is responsible for the UV-to-violet shift between the two SWS1 opsins. The novel violet-sensitive SWS1B might compensate, at least in part, for the ancestral loss of blue opsins in snakes. The presence of two SWS1 provides a potential substrate for trichromatic or even tetrachromatic vision in this lineage. Finally, we find that the unique expansion of the opsin toolbox in *Helicops* was accompanied by morphological specializations of the retina such as a higher density of photoreceptors and short-wavelength sensitive cones in the ventral retina, possibly reflecting the freshwater habitats of this lineage.

### Both UV- and violet-sensitive SWS1 opsins are expressed in the Helicops retinas

Micro-spectrophotometry showed that both UV and violet single cones (together with long wavelength cones and rods) can be simultaneously present in *Helicops* retina. The spectral peaks of the UV-sensitive and the violet-sensitive SWS1 cones measured by MSP matched the theoretical predictions based on tuning sites and directly measured by *in vitro* expression of the reconstructed SWS1 opsins. The extent of sequence divergence in exonic and intronic regions and the patterns of phylogenetic clustering of the two *sws1* variants indicate that these represent distinct paralogous genes that experienced different selective pressures, as evidenced by the d_N_/d_S_.

The simultaneous expression of a UV and a violet SWS1 opsin is unprecedented in any vertebrate. Genome mining suggested that *sws1* duplications might have independently occurred in some fish lineages (Minamoto and Shimizu 2005; Lagman et al. 2013; Musilova et al. 2019), though it is not known whether such duplicates represent functional genes simultaneously expressed in a retina and sensitive to distinct wavelengths. In *Helicops* snakes, the two distinct SWS1 opsins might have evolved as a secondary adaptation to freshwater aquatic habitats, reducing the spectral gap between the ancestral UV-sensitive cones and the yellow-red sensitive LWS cones. An intriguing possibility is that the pathways beginning with UV and violet cone excitation might lead to post-receptoral opponent responses, allowing color discrimination in the very short waveband, an exciting venue for future behavioral and physiological studies. Alternatively, the two SWS1 opsins might have evolved completely distinct roles not involved in color vision, but specialized in other functions such as polarization vision, as proposed for UV cones in other vertebrates (Bennett and Cuthill 1994; Hawryshyn 2010).

A recent study on marine snakes of the genus *Hydrophis* (Simões et al. 2020) analyzed opsin genomic sequences finding allelic polymorphisms across species at a main SWS1 tuning site. The authors speculated that heterozygous individuals might express both alleles, and gain an advantage over homozygous ones in a similar fashion to that enjoyed by heterozygote females of New World monkeys. Based on known tuning sites, Simões et al. (2020)’s study provided approximate predictions for the spectral sensitivity of each allele. However, evidence that two SWS1 opsins are simultaneously expressed in these snakes is still lacking. In addition, the proposed fitness advantage of heterozygotes would only be present if both alleles were expressed not only simultaneously but in codominant fashion in distinct photoreceptors.

The marine elapids of Simões et al. (2020) and the freshwater *Helicops* dipsadids studied here might have followed different evolutionary routes during their invasion of aquatic environments, leading to the expansion of the visual opsin repertoire, a diversity of adaptive routes previously documented in primates. Mammal ancestors passed through a nocturnal bottleneck that resulted in the loss of visual structures and two cone opsins sensitive to the core region of the visible spectral, SWS2 and RH2 (Davies et al. 2012; Gerkema et al. 2013). Some primates are unique among mammals in displaying trichromatic color vision based on SWS1, MWS, and LWS opsins, a gain achieved through two distinct mechanisms in different lineages (Carvalho et al. 2017). In Old World monkeys, an *lws* duplication followed by substitutions at spectral tuning sites resulted in spectrally distinct opsins (MWS and LWS) conferring trichromacy in both males and females (Nathans et al. 1986; Hunt et al. 1998; Dulai et al. 1999). In most New World monkeys, a high degree of polymorphism at the spectral tuning sites of a single X-linked *lws* locus give rise to multiple pigments with distinct *λ*_max_, resulting in a conditional form of trichromacy in heterozygous females (Mollon et al. 1984; Carvalho et al. 2017). Worth noting, substitutions at the same residues are responsible for the spectral shift of the MWS and LWS opsins in both primate lineages. In snakes, the fact that the same opsin site 86 was involved in SWS1 tuning in two independent lineages, might represent a remarkable example of convergent evolution in which distinct lineages independently evolved similar phenotypes through the recruitment of substitutions at the same residue. The invasion of aquatic environments by a terrestrial vertebrate is probably at the origin of convergent visual phenotypes in these two independent snake lineages but the presence of mutations at the same site underlying such convergence points to mutational biases or genetic constraints in this opsin (Losos 2011; Stern 2013), an exciting avenue for future research.

### The basis for spectral tuning of UV and violet photopigments in Helicops is a single substitution

The SWS1A and SWS1B opsins of *Helicops* have different amino acids at residue 86, the main SWS1 spectral tuning site (Cowing et al. 2002; Fasick et al. 2002; Parry et al. 2004). In SWS1A, residue Phe86 points to sensitivity in the UV range, while Val86 in SWS1B suggests sensitivity in the violet range (Parry et al. 2004). These predictions were confirmed by our *in vitro* expression of the *sws1a* and *sws1b* genes of *H. modestus*. We showed that both proteins transcribed are functional and have distinct phenotypes, SWS1A with *λ*_max_ at 364nm, and SWS1B, with *λ*_max_ at 417nm, values remarkably close to the absorbances that we measured directly in individual photoreceptors with MSP. Moreover, using site-directed mutagenesis, we confirmed, for the first time, the functional role of the single amino acid substitution at residue 86 in generating the wide spectral shift of UV and violet photopigments in this lineage of snakes. While precise MSP estimates of the *λ*_max_ of these two SWS1 opsins will require a larger dataset, the substantial *λ*_max_ shift of the UV-SWS1A to longer wavelength (395nm) recorded by preliminary MSP measurements in *H. angulatus* is likely to be caused by additional amino acid residues, such as Val93, a spectral tuning site that differed between *H. angulatus* and other *Helicops* species. In the majority of snakes investigated so far, residue 86 is thought to predict well the spectral peak of SWS1 opsins (Davies et al. 2009; Schott et al. 2016; Bhattacharyya et al. 2017; Simões et al. 2020). However, in the viperid, *Agkistrodon contortrix*, the UV prediction based on this site (Phe86) does not correspond to the *λ*_max_ in the violet range (416nm) recorded by MSP (Gower et al. 2019). This discrepancy points to the relevance of additional residues in the *λ*_max_ of snakes SWS1, in particular 93, a candidate site for future site-directed mutagenesis assays.

The SWS1 is the only vertebrate visual opsin class that can reach a spectral sensitivity peak in the UV range, the consequence of the absence of protonation in the chromophore Schiff base in the dark state (Fahmy and Sakmar 1993). In the other visual opsins, including violet SWS1 photopigments, the Schiff base of the 11-cis-retinal chromophore is protonated, with a highly conserved glutamate counterion (E113) acting on the proton stabilization (Nathans 1990). Site-directed mutagenesis assays demonstrated that substitutions at site 86 from various amino acids (Tyr, Ser, Leu, Val) characteristic of violet-sensitive SWS1 opsins in mammals, and now snakes, to Phe86, cause the loss of electrostatic stabilization of the protonation, shifting the absorption peak of the photopigment to the UV range (Cowing et al. 2002; Parry et al. 2004; Yokoyama et al. 2005; Carvalho et al. 2012). On the other hand, the non-polar residues Val86, Met86, and Leu86, despite being naturally found in violet opsins of guinea pig, primates and amphibians, respectively, were not able to generate any spectral change in UV opsins of fish (Cowing et al. 2002; Hunt et al. 2004; Parry et al. 2004). In our study, the substitution Phe86Val introduced in the UV opsin of *H. modestus*, unexpectedly generated a considerable shift to the violet, indicating stabilization of the protonation of the Schiff base. This is probably achieved by the interaction of residue Val86 with other spectral tuning sites, such as the non-polar residue Met93 (Shi et al. 2001). However, we observed considerable noise in the mutant Phe86Val opsin curve possibly caused by a low yield due to reduced stability of the generated opsin (Hauser et al. 2014), or a possible equilibrium between unprotonated (UV) and stabilized protonated (violet) Schiff base linkage of the reconstituted opsin mutants. The resulting curve did not optimally fit a typical opsin template which led to only an approximate estimate of the spectral peak, around 384nm.

### Distinct patterns of selection on sws1a and sws1b opsins

We detected a remarkable difference in the evolutionary rates of the two *Helicops sws1* opsin genes. Clade models indicate positive selection in the ancestral branch leading to *Helicops* sequences (ω > 1) at the origin of the suggested *sws1* duplication, pointing to accelerated rates of evolution that eventually resulted in the fixation of novel opsin phenotypes. However, the analyses also indicated ω << 1 on the ancestral UV-sensitive *sws1a*, suggesting that this opsin paralog is typically evolving under strong purifying selection to maintain receptor functionality and characteristic sensitivity. A signal of strong purifying selection on *sws1* opsins was also found in other snakes, with ω values that do not exceed 0.2 (Simões et al. 2016; Hauzman et al. 2017). On the other hand, clade models indicated values of d_N_/d_S_ ∼1 for the second *sws1* paralog, the novel violet-sensitive opsin *sws1b*. Different d_N_/d_S_ trajectories of young duplicated genes are to be expected in a neo-functionalization scenario, where an early phase of strong directional selection at few sites will work in conjunction with strong purifying selection on the rest of the gene. After this transient stage establishing the new paralog in the population, purifying selection will act ubiquitously on this gene, as in the ancestral copy, resulting in a gradual return to pre-duplication evolutionary rates.

Pegueroles et al. (2013) studied the dynamics of evolutionary rates in recent paralogs by examining 404 duplicated gene copies generated at different times during rodent evolution. The authors found that asymmetry in ω estimates between duplicates is common, with the novel daughter copy typically evolving more rapidly. In their comprehensive study, the observed acceleration was limited to a relatively short period of time following the duplication event, gradually decreasing and being completely erased after approximately 40.5 My. While our data do not allow reliable estimates of divergence time for the two *sws1* paralogs, species trees indicate that the genus *Helicops* diverged about 13.9 My ago from the other members of its clade (Zaher et al. 2019). This places the origin of the *sws1* paralogs in a time window consistent with Peguelores et al. (2003)’s phase of strong directional selection at one or few sites and strong purifying selection on the rest of the gene, leading to the observed d_N_/d_S_ ∼1, as the signatures of the brief positive selection spike are gradually eroded, after fixation of the new paralog.

### Ecological implications of a broadened sensitivity at short wavelengths

The fate of a duplicated gene is largely determined by its potential to evolve new functions (Ohno 1970; Walsh 2003; Gojobori and Innan 2009). In fish, where opsin gene duplications are not uncommon, paralogs of opsins sensitive to the central portion of the spectrum (i.e. blue and green *sws2* and *rh2* opsins) are much more common than duplications involving opsins sensitive to either end of the visible light spectrum (*sws1* and *lws*) (Gojobori and Innan 2009). Duplications of the *sws1* gene are rare and, to the best of our knowledge, have been suggested only in few fish species (Minamoto and Shimizu 2005; Musilova et al. 2019). However, no study has demonstrated the simultaneous expression of more than one *sws1* variant in a retina of any vertebrate. In *Helicops* snakes, the gain of a new violet opsin in addition to the UV opsin might have been selected upon to partially compensate for the ancestral loss of the SWS2 opsin. The additional violet opsin of *Helicops* snakes fills a spectral gap between the UV and LWS opsins, enabling a broader coverage of the visible light spectrum available. Generally, freshwater environments have less light at short wavelengths, especially in the UV range, due to suspended organic matter and rapid absorption of the shortest wavelengths with depth (Loew and Lythgoe 1985; Lythgoe and Partridge 1989). The life of a *Helicops* snake is at the interface of two drastically different visual worlds, terrestrial and aquatic. The unique adaptations of its visual system reflect this duality: underwater, a violet pigment may enable efficient photon capture at short wavelengths and, together with LWS cones, provides the potential for color vision. Above water, UV light is abundant and the snake can take full advantage of the broader light space available by employing the additional UV photoreceptor, and potentially gaining color discrimination at the shortest wavelengths.

### Retinal structure and the visual ecology of Helicops snakes

*Helicops* snakes are predominantly nocturnal (Martins and Oliveira 1998; De Aguiar and Di-Bernardo 2004). However, unexpectedly, their retinal structure is similar to that of diurnal snakes, with a low density of photoreceptors, absence of typical rods (Walls 1942; Underwood 1967; Wong 1989; Sillman et al. 1997; Hart et al. 2012; Hauzman et al. 2014; Hauzman et al. 2017), and the presence of transmuted cone-like rods. The low density of photoreceptors points to lower light sensitivity compared to nocturnal snakes, with duplex retinas and high density of typical rods (Walls 1942; Sillman et al. 1999; Sillman et al. 2001; Hauzman et al. 2017). A similar apparent incongruence was described in the nocturnal dipsadid *Thamnodynastes*, with a typical diurnal retina (Hauzman 2014). It was suggested that a shift to nocturnal activity in *Thamnodynastes* is associated with hunting anurans (Torello-Viera and Marques 2017), while the diurnal retinal pattern may be due to phylogenetic inertia, so that the change in the circadian rhythm might have not been accompanied (yet) by changes in retinal morphology. More work on the diel activity patterns of *Helicops* is needed to evaluate whether similar selective forces are at the origin of its typically diurnal retina.

So far, only a handful of studies have described retinal topography in snakes (Wong 1989; Hart et al. 2012; Hauzman et al. 2014; Hauzman et al. 2018). In dipsadid snakes, a visual streak was found in the arboreal *Philodryas olfersii* (Hauzman et al. 2014), a specialization that might enable the view of potential predators and prey located at the same level of the snake. On the other hand, in the closely related terrestrial species, *Philodryas patagoniensis*, a ventral anisotropic *area centralis* was described, which would enable detection of predators from above (Hauzman et al. 2014). In *Helicops modestus* we observed higher density of photoreceptors and SWS cones in the ventral retina, as in *P. patagoniensis*, likely resulting in higher resolution of the upper visual field, in addition to a higher photoreceptor absorption of short-wavelength photons from downwelling light. In mice, a ventral retina dominated by UV cones (Szél et al. 1992; Röhlich et al. 1994) might improve the view of potential aerial predators against a UV-rich clear sky (Szatko et al. 2020). Behavioral studies revealed chromatic discrimination in the superior visual field of the murine (Jacobs et al. 2004; Denman et al. 2018), and opponent chromatic channels are provided by inputs from UV cones and rods (Joesch and Meister 2016), a functional organization associated with an ability to perceive contrast from dark objects against a clear sky background under mesopic conditions, in which cones and rods are functional, and mice are active (Joesch and Meister 2016; Szatko et al. 2020). In *Helicops*, it would be valuable to investigate whether similar selective pressures led to the observed retinal organization, enhancing the ability to perceive dark objects against a brighter background such as aerial predators, and allowing a fast escape dive underwater. Indeed, a wide variety of birds prey on aquatic and terrestrial snakes (Guthrie 1932; Mushinsky and Miller 1993), including *Helicops* (Franz et al. 2007). However, one must exert caution when interpreting retinal organization at the photoreceptor level given the complexity of visual processing pathways within the retina and beyond (Baden et al. 2020). A second possible benefit conferred by a ventral *area centralis*, might be a diversification of the snake’s hunting strategies. This could be achieved through higher acuity or improved chromatic discrimination in the upper visual field, facilitating the detection of anurans on aquatic plants above the snake. Further investigations on retinal circuitry, including photoreceptor connectivity to bipolar and ganglion cells, and visually-guided behaviors will be required to fully understand the adaptive link between the retinal organization, visual function, and the life history of *Helicops*.

This study shows, for the first time in a vertebrate species, the presence of two distinct SWS1 opsins, one UV-sensitive and the other violet-sensitive, simultaneously expressed in the retinas of a lineage of the aquatic *Helicops* snakes. We demonstrated that the shift in spectral sensitivity between the UV- and violet-sensitive opsins is largely due to a single amino acid substitution at a known tuning site. Additional unique features of the visual system of this aquatic snake lineage described here suggest higher acuity in behaviorally relevant directions and possibly higher chromatic discrimination in the complex light environments of freshwater habitats, in addition to specialized short wavelength photon capture from the upper visual field. Our study highlights the extraordinary potential for rapid evolution of vertebrate visual opsins and the remarkable diversity of visual adaptations characterizing the ophidian tree of life.

## Material and Methods

### Sample Information

Snakes (n=13) were euthanized with a lethal injection of 100 mg/kg of sodium thiopental (Thionembutal). All procedures were in accordance with ethical principles of animal management and experimentation of the Brazilian Animal Experiment College (COBEA), and approved by the Ethics Committee of Animal Research of the Butantan Institute, São Paulo, Brazil (777/10; 4479020217) and the Psychology Institute, University of São Paulo, Brazil (9635070717). The voucher specimens were fixed and deposited in the Herpetological Collection of the Butantan Institute, São Paulo, or in the Herpetological Collection of the Zoological Museum of the University of São Paulo (supplementary table S10).

### RNA extraction, PCR, cloning, and sequencing

The eyes of single specimens of four *Helicops species, H. modestus*, *H. infrataeniatus*, *H. carinicaudus*, and *H. leopardinus*, and *Hydrops caesurus* were enucleated and preserved in RNAlater® (Life Technologies, Carlsbad, California) at 4°C. Total RNA was extracted from homogenized retinas using the RNase Mini Kit (Qiagen GmbH, Hilden, Germany) according to the manufacturer’s instructions and preserved at −80°C. Total RNA was diluted 10 fold and mRNA was converted to complementary DNA (cDNA) using 500 ng of oligo-dT primer and the reverse transcriptase MultiScribe™ (Applied Biosystems, Foster City, California), following the manufacturer’s protocol. Polymerase chain reactions (PCRs) were performed to amplify the *sws1*, *rh1*, and *lws* opsin genes, using High Fidelity Platinum Taq Polymerase, in 50 μL reactions, with 10x High Fidelity Buffer, MgCl_2_, 10mM GeneAmp dNTPs (Life Technologies, Carlsbad, California), and 20 μM primers (supplementary table S11). The PCR conditions were: i) denaturation at 94°C for 1 min; ii) 37 cycles at 94°C for 15 s, annealing temperature for 30 s, and extension temperature at 72°C for 30 s; iii) 10 min of extension temperature. The PCR products were visualized by electrophoresis in 1.0% agarose gel, and purified using the Illustra GFX^TM^ PCR DNA and Gel Band purification kit (GE Healthcare, Little Chalfont, Buckinghamshire, UK). The purified PCR products were directly sequenced in both directions, using BigDye® Terminator v3.1 Cycle Sequencing Kit (Applied Biosystems) and a 3500 Applied Biosystems Sequencer. Electropherograms were visualized with BioEdit v7.2.5 (Hall 1999). For the *sws1* opsin gene, PCR products with ∼1000 bp were cloned into plasmid vectors using TA Cloning® kit (Life Technologies, Carlsbad, California). Colonies were screened by blue/white selection, and multiple selected positive clones were isolated, purified via spin columns (Qiagen GmbH, Hilden, Germany), and sequenced using T7 and M13 primers.

### Phylogenetic analysis

Resulting sequences were visualized and aligned with Geneious v.9.1.3 (GeneMatters Corp.), using the iterative method of global pairwise alignment (MUSCLE and ClustalW) (Thompson et al. 1994; Edgar 2004). The alignments of each opsin gene, *lws*, *rh1*, and *sws1*, included the sequences generated in this study and other snake opsin gene sequences obtained from GenBank (supplementary table S12). Maximum Likelihood (ML) reconstructions were performed on codon-match nucleotide alignments, using Garli v2.0 (Bazinet et al. 2014). Partition-Finder v.1.1.1 (Lanfear et al. 2012) was used to determine the best fit-models: models TrNef+I+G, TVM+I+G, and GTR+G were the best fit-models for codon positions 1, 2, and 3 of the *sws1* gene, and were used in ML reconstruction. Models TIMef+I+G, TVM+I+G, and TrN+G were used for positions 1, 2, and 3 of the *rh1* gene, and for the *lws* gene, models TIM+I+G, TVM+I+G, and K81uf+G were used for position 1, 2 and 3, respectively. Statistical support was estimated by non-parametric bootstrap (Felsenstein 1985), with 1000 pseudo replications.

### Genomic analysis of the sws1 opsin

We amplified and sequenced the exon 1 of the *sws1* gene from genomic DNA to search for variants in single individuals from six *Helicops* species (*H. modestus*, *H. angulatus*, *H. hagmanni*, *H. polylepis*, *H. leopardinus*, and *H. gomesi*), two *Hydrops* (*H. triangularis* and *H. martii*), *Pseudoeryx plicatilis*, and two non-Hydropsini aquatic colubris, *Sordellina punctata* and *Hydrodynastes gigas*. Tissue samples were donated by the Vertebrate Tissue Collection from the Department of Zoology, University of São Paulo (supplementary table S13). DNA was extracted from liver tissues using standard protocols (Puregene DNA®, Gentra System), and PCRs were performed as described above (supplementary table S11). PCR products were visualized in agarose gel 1%, purified and sequenced as described previously. In addition, we amplified by PCR the intron 1 of single individuals from the same six *Helicops* species, using combinations of specific primers pairs (supplementary table S11). The amplified DNA fragments were purified from 1.5% UltraPure™ low melting point agarose gel (Invitrogen™). Sequencing was performed in both directions as described above. In two species, *H. modestus* and *H. leopardinus*, two size-divergent introns were identified. After a first round of sequencing analysis, additional specific primers were designed to separately amplify and sequence the two introns.

### Estimates of opsin spectral tuning

We predicted the wavelength of maximum absorption (*λ*_max_) of the visual pigments expressed in the retinas of the newly sequenced species. Estimates were made on known spectral tuning amino acid sites, using the conventional numbering based on the bovine rhodopsin sequence. For the LWS opsin, residues S164, H181, Y261, T269 and T292 are known to generate a *λ*_max_ at ∼560 nm (Yokoyama and Radlwimmer 1998), and the substitutions S164A, H181Y, Y261F, T269A and A292S to cause downward shifts of 7, 28, 8, 15 and 27 nm, respectively (Yokoyama and Radlwimmer 2001). For the rhodopsin photopigment, we analyzed 26 amino acid sites: 83, 90, 96, 102, 113, 114, 118, 122, 124, 132, 164, 183, 194, 195, 207, 208, 211, 253, 261, 265, 269, 289, 292, 295, 299, and 317 (Chan et al. 1992; Kawamura et al. 1999; Yokoyama et al. 1999; Fasick and Robinson 2000; Yokoyama 2000; Hunt et al. 2001; Janz and Farrens 2001; Yokoyama 2008; Yokoyama et al. 2008). For the SWS1 opsin we analyzed the spectral tuning sites 46, 49, 52, 86, 90, 93, 97, 113, 114, 116, 118, 265 (Shi et al. 2001; Yokoyama et al. 2006; Yokoyama 2008), and the absorption peak at UV range was determined based on the presence of the amino acid Phe86 (Cowing et al. 2002; Fasick et al. 2002).

### *In vitro* expression of the SWS1 opsins and site-directed mutagenesis

Two distinct *sws1* opsin genes, here named *sws1a* and *sws1b*, were identified in retinas of *Helicops* snakes. To investigate the functional properties of these opsins, we expressed *in vitro* both *sws1* copies from the species *H. modestus*. The *sws1a* and *sws1b* coding sequences along with a C-terminal nine-amino acid epitope tag for the 1D4 antibody were synthesized (GenOne Biotechnologies, Rio de Janeiro, Brazil), and inserted into the p1D4-hrGFP II expression vector (Morrow and Chang 2010). Expression vectors were transiently transfected into cultured HEK293T cells (ATCC CRL-11268), using Lipofectamine 2000 (Invitrogen; 8 μg of DNA per 10-cm plate), and harvested after 48 h, with Harvesting Buffer (50mM HEPES ph 6.6, 140mM NaCl, 3mM MgCl_2_). Bovine rhodopsin was transfected along with the *sws1* genes for control, following the same protocols. Opsins were regenerated with the 11-cis retinal chromophore, generously provided by Dr Rosalie Crouch (Medical University of South Carolina), solubilized in 1% n-dodecyl-β-D-maltopyranoside detergent (DM) with 20% (w/v) glycerol, and purified with the 1D4 monoclonal antibody (University of British Columbia 95-062, lot 1017) (Molday and MacKenzie 1983), as previously described (Morrow and Chang 2010; Morrow and Chang 2015). The opsins were purified in HEPES buffers containing glycerol (van Hazel et al. 2013). Site-directed mutagenesis were performed via PCR following the QuikChange site-directed mutagenesis protocol (Agilent) and using PfuUltra II Fusion HS DNA Polymerase (Agilent), to generate mutants: SWS1A-Phe86Val and SWS1B-Val86Phe. Mutagenesis primers were designed with 37 nucleotides identical to each gene flanking the mutant nucleotide. Mutants were confirmed by double-stranded sequencing, and inserted into p1D4-hrGFP II expression vectors, which were used for transfection into cultured HEK293T cells, along with bovine rhodopsin as control, as described above. The UV-visible absorption spectra of wild and mutant purified visual pigments were recorded using a Cary 4000 double beam spectrophotometer (Agilent). The photopigments were photo-excited with light from a fiber optic lamp (Dolan-Jenner, Boxborough, MA, USA) for 60 s at 25°C. To estimate the wavelength of maximum absorption, the dark absorbance spectra were baseline-corrected and fit to Govardovskii et al. (2020)’s template curves for A1 visual pigments.

### Statistical analysis of molecular evolution

We investigated the presence and type of selection acting on the *sws1* opsin genes by applying a codon-based method, using the codeml program from PAML package v.4.7 (Yang 2007), in a Dipsadidae alignment with 897 bp (species listed in supplementary table S12), using the gene tree (fig. 1a) and a species tree (Pyron et al. 2013). The ratio of nonsynonymous (d_N_) to synonymous (d_S_) substitutions (ω), indicates the type and magnitude of selection, where ω < 1 indicates purifying selection, ω ∼ 1 indicates neutral evolution, and ω > 1 indicates positive selection (Yang and Nielsen 1998; Yang 2007), and were estimated using random-sites models, branch-site models, and clade (CmC) models (Bielawski and Yang 2004; Zhang 2005; Yang 2007; Weadick and Chang 2012). Likelihood ratio tests (LRTs) were used to compare competing models of evolution and were computed as 2log likelihood difference between two models and tested against the χ2 distribution, where the degrees of freedom equal the difference between the numbers of parameters in the two nested models (Yang 2007). Models were also compared according to their likelihood using Akaike’s information criterion (AIC) (Akaike 1974). Random-sites models allow ω to vary among codon sites and to detect sites potentially under positive selection. Random-sites, M0, M1a, M2a, M2a_rel, M3, M7, M8a, and M8 (Yang 2007; Weadick and Chang 2012), were used to determine the overall selective patterns and to test for heterogeneous selection pressure among codon sites across all branches of the tree. LRT comparisons between random site models were used to test for variation in ω among sites (M3 vs. M0) and the presence and proportion of positively selected sites (M2a vs. M1a, M8 vs. M7, and M8 vs. M8a) (Yang et al. 2000; Yang 2007). When LRTs were significant for positive selection, we used Bayes Empirical Bayes (BEB) to estimate posterior probabilities for site classes and identify amino acid sites under positive selection (Yang 2007). Branch-site and CmC models allow to test for divergence along specific branches of the tree appointing them as “foreground” and compare their ω rates with that estimated for the “background” branches (Zhang 2005; Yang 2007). Branch-site models were used to analyze whether specific clades of interest have experienced positive selection on any codon site, by isolating in independent foregrounds: i) the aquatic clade (Hydropsini snakes: *Helicops* species and *Hydrops caesurus*); ii) the *Helicops* clade; iii) the *sws1a* clade; and iv) the *sws1b* clade. We implemented Model A (Model = 2, NSsites = 2) as an extension of the site-specific “neutral” model (M1) (Nielsen and Yang 1998). The null models are the same as for Model A but with ω_2_ fixed at 1 for foreground branches. The proportions p_0_ and p_1_, as well as the ratio ω_2_, were estimated by maximum likelihood (Nielsen and Yang 1998; Yang et al. 2000). We used CmC models to test the hypothesis of divergent evolution in specific branches and clades. CmC allows two classes of sites across the tree to evolve conservatively (0 < ω < 1) and neutrally (ω = 1), while a third site class is free to evolve differently. The CmC null model, M2a_rel, does not allow ω to diverge in the foreground clade (Weadick and Chang 2012). We applied two-partition models to investigate the patterns of selection acting on the ancestral branches leading to i) aquatic snakes, ii) *Helicops* snakes, iii) *sws1a*, and iv) *sws1b*, and at the same respective clades, as independent foregrounds against the background. In a four-partition scheme, we isolated as foregrounds i) the *Helicops* ancestral branches, ii) the *sws1a* clade, and iii) the *sws1b* clade, against the background.

### Microspectrophotometry

The photoreceptors spectral sensitivities of *H. angulatus* (n=1) and *H. leopardinus* (n=3) were investigated with microspectrophotometry. Snakes were dark adapted for one hour before being euthanized. Eyes were enucleated and the cornea removed under dim red light, and eyecups were immersed overnight in calcium-free phosphate-buffered saline (PBS) buffer (Sigma-Aldrich) with sucrose solution (6.0%). Preparations were particularly challenging due to a thick and sticky pigment epithelium that required careful manipulation, often causing detachment of the extremely delicate and short outer segments of cones. Small pieces of retina detached from the pigment epithelium were placed on glass cover slips in PBS solution and gently macerated using razor blades. The preparation was covered with a second cover slip and sealed with high-vacuum silicone grease (Dow Corning). Spectral absorbance was measured with a computer-controlled single-beam microspectrophotometer fitted with quartz optics and a 100W quartz-halogen lamp. Baseline records were taken by averaging a scan from 750 nm to 350 nm and a second in the opposite direction. Records of selected photoreceptors were obtained by subtracting a baseline record obtained in an area free of cells from a beam scan through the photoreceptor. Maximum absorption peaks were obtained by fitting A1 templates (Govardovskii et al. 2000) to the smoothed, normalized absorbance spectra.

### Ocular media transmittance

We measured lens and spectacle transmittance of *H. leopardinus* (n=2) using the methods described by Lind et al. (2013) and Yovanovich et al. (2019). The samples were placed on top of a black plastic disc with a 1mm pinhole inside a custom-made matte black plastic cylinder (12mm diameter×10mm height) with a circular (5mm diameter) fused silica window in the bottom, filled with PBS. We used an HPX-2000 Xenon lamp (Ocean Optics, Dunedin, FL, USA) to illuminate the samples via a 50μm light guide (Ocean Optics) and collected transmitted light using a 1000μm guide connected to a Maya2000 spectroradiometer controlled by SPECTRASUITE v.4.1 software (Ocean Optics). The guides were aligned with the container in a microbench system (LINOS, Munich, Germany). The reference measurement was taken from the container filled with PBS. We smoothed the curves using an 11-point running average and normalized to the highest value within the range 300–700nm. From these data, we determined *λ*_T50_ as the wavelength at which the light transmittance was 50% of the maximum.

### Retinal morphology

We investigated the retinal structure and photoreceptor types using immunohistochemistry in retinal section of *H. modestus* (n=1) and *H. carinicaudus* (n=1). After eye enucleation, the cornea and lenses were removed and the eyecups were fixed in 4% paraformaldehyde (PFA) diluted in PBS, for 3 hours. The eyecups were cryoprotected with 30% sucrose solution for 48 hours, and sectioned at cryostat (CM1100 Leica, Nussloch, Germany) at 12-μm thickness. Sections were pre-incubated for 1 hour in 10% normal goat serum (NGS) or normal donkey serum (NDS) (Sigma-Aldrich), diluted in PBS with 0.3% Triton X-100 and incubated overnight at room temperature with anti-opsin antibodies: rabbit anti-SWS1 (Chemicon International; AB5407; 1:200), goat anti-SWS1 (Santa Cruz Biotechnology; sc-14363; 1:200), rabbit anti-LWS (Chemicon International; AB5405; 1:300), and mouse anti-rhodopsin RET-P1 (EMD Millipore, MAB5316; 1:200), diluted in PBS with 0.3% Triton X-100. Three mixtures of primary antibodies were used for double immunofluorescence labeling and analysis of opsins coexpression: i) rabbit anti-SWS1 (AB5407) combined with mouse anti-rhodopsin (MAB5316); ii) rabbit anti-LWS (AB5405) combined with mouse anti-rhodopsin (MAB5316); and iii) goat anti-SWS1 (sc-14363) combined with rabbit anti-LWS (AB5405). The specificity of the antibodies for snakes were described previously (Bhattacharyya et al. 2017; Hauzman et al. 2017). The sections were washed in PBS with 0.3% Triton X-100, and incubated for 2 hours with the following combinations of secondary antibodies: i) fluorescein (FITC)-conjugated goat anti-rabbit with cyanine (CY3)-conjugated goat anti-mouse; and ii) Alexa Fluor® 488-conjugated donkey anti-rabbit with tetramethylrhodamine (TRITC)-conjugated donkey anti-goat (immunoglobulin G, whole molecules; 1:200; Jackson Immunoresearch Laboratories). No labeling was observed in negative control sections, with the omission of the primary antibodies. Sections were washed in PBS and mounted with Vectashield with 4,6-diamidino-2-phenylindole (DAPI; Vector Laboratories), coverslipped, and observed under a fluorescent microscope (Leica DM5500B). A Zeiss LSM 880 confocal microscope in Airyscan mode was used for super resolution imaging, with a Zeiss x63/1.4 oil immersion objective (Carl Zeiss, Germany), with a combination of filters for visualizing the fluorophores used: for DAPI staining, excitation wavelength at 405 nm and emission at 455 nm, for Alexa Fluor® 488 and FITC-conjugated antibodies, excitation at 488 nm and emission at 516 nm, and for TRITC and CY3-conjugated antibodies, excitation at 543 nm and emission at 594 nm. The images were analyzed using Zen Blue (Zeiss), and labeling colors of each filter used were changed using the same software, to conveniently differentiate the opsin types (LWS, SWS1, and RH1), irrespective of the dye used.

### Density and topographic distribution of all photoreceptors and of SWS1 cones

We investigated density and distribution of photoreceptors and SWS1 cones in wholemount retinas of *H. modestus* (n=3). After eyes enucleation, the cornea and lenses were removed and a small radial incision was made in the dorsal region for later orientation. The eyecups were fixed in 4% PFA diluted in PBS for 3 hours. Retinas were carefully dissected from the eyecup, and when the pigment epithelium was not easily separated from the retina, we treated with bleaching solution of 10% hydrogen peroxide diluted in PBS, overnight at room temperature. Free-floating retinas were pre-incubated for 1 h in 10% NGS (Sigma-Aldrich), incubated with anti-SWS1 opsin antibody (1:100) for 72 hours, and with secondary antibody, goat anti-rabbit immunoglobulin G (1:200), conjugated with FITC. Strategic cuts were made in the retinas to allow them to be flattened. Slides were mounted with Vectashield and coverslipped. To analyze the density and distribution of cells we used a systematic random sampling and the fractionator principle (West et al. 1991), modified for retinal wholemounts (Coimbra et al. 2009). The total number of cells were estimated based on area sampling fraction (asf), i.e. the ratio between the counting frame and the sampling grid, according to the algorithm: N_total_ = ΣQ x 1/asf, where ΣQ is the sum of total neurons counted (West et al. 1991). The degree of accuracy was calculated using Scheaffer coefficients of error. The retinas were outlined and sampling grids with approximately 200 fields were placed in a random and uniform distribution, covering the whole retinal area, using a microscope (Leica DM5500B), equipped with a motorized stage and connected to a computer running Stereo Investigator software (MicroBrightField). Cells were counted using a x40/0.8 objective, if they were fully inserted within the counting frame or if touched the acceptance lines, without touching the rejection lines (Gundersen 1977). The cell densities per square millimeter at each counting frame was used to elaborate the topographic maps, with the software OriginPro 8.1. The images were processed using the software Adobe Photoshop CS3 (Adobe Systems, Inc.).

## Acknowledgments

We thank Felipe Franco Curcio, Felipe Gobbi Grazziotin and Giuseppe Puorto for providing the snakes, Valdir Germano, Lucas Amâncio Neves, and Bruno Rocha for assisting on the animals’ access, Karina Banci and Natalia Torello-Viera for providing preliminary data on snake’s daily activity patterns. We thank Waldir Caldeira for obtaining confocal images and Andrea Vieira de Souza for crucial assistance with Sanger sequencing. We are also indebted to Inacio L.M. Junqueira de Azevedo for access to facilities at Butantan Institute and to Miguel Trefaut Rodrigues for providing tissue samples and for valuable discussions. Our gratitude also goes to Professor Ellis Loew for generously lending his microspectrophotometer for this study and for helpful advice on retinal preparation. Sequences data generated in this study will be deposited in the NCBI database, and accession numbers will be provided upon acceptance. This work was supported by The São Paulo Research Foundation (FAPESP) (grants numbers: 2014/25743-9 and 2018/09321-8 to EH, 2014/26818-2 to DFV, 2018/11502-0 to MERP, 2018/13910-9 to JHT, 2015/14857-6 to CAMY, 2013/07467-1 to PFC); from the Brazilian Coordination for the Improvement of Higher Education Personnel (CAPES) (grant number 08307/03/2013 to PFC); and from the Brazilian National Research Council (CNPq) (grant number 309409/2015-2 to DFV).

## Data Availability

The data underlying this article will be available in the GenBank Nucleotide Database (to be provided).

## Author Contributions

E.H., D.F.V., and B.S.W.C conceived the study and obtained funding. E.H., M.E.R.P, N.B, J.H.T, C.A.M.Y., and P.F.C. performed research. E.H., M.E.R.P., N.B., J.H.T., C.A.M.Y., D.F.V., and B.S.W.C. analyzed data. E.H. and M.E.R.P wrote the paper. N.B., J.H.T., C.A.M.Y., D.F.V., and B.S.W.C. contributed comments to the paper.

